# Control of Cardiac Mitochondrial Fuel Selection by Calcium

**DOI:** 10.1101/198895

**Authors:** Edith Jones, Sunil M. Kandel, Santosh K. Dasika, Neda Nourabadi, Françoise Van den Bergh, Hyo Sub Choi, Ali Haidar, Ranjan K. Dash, Daniel A. Beard

## Abstract

Calcium ion concentration modulates the function of pyruvate dehydrogenase, isocitrate dehydrogenase, and α-ketoglutarate dehydrogenase. Previous studies have shown that despite its ability to affect the function of these dehydrogenases, [Ca^2+^] does not substantially alter mitochondrial ATP synthesis in vitro under physiological substrate conditions. We hypothesize that, rather than contributing to respiratory control, [Ca^2+^] governs fuel selection. Specifically, cardiac mitochondria are able to use different primary carbon substrates to synthesize ATP aerobically. To determine if and how [Ca^2+^] affects the relative use of carbohydrates versus fatty acids we measured oxygen consumption and tricarboxylic acid cycle intermediate concentrations in suspensions of cardiac mitochondria with different combinations of pyruvate and palmitoyl-L-carnitine in the media at various [Ca^2+^] and ADP infusion rates. Results reveal that when both fatty acid and carbohydrate substrates are available, fuel selection is sensitive to both calcium and ATP synthesis rate. When no Ca^2+^ is added under low ATP-demand conditions, β-oxidation provides roughly half of acetyl-CoA for the citrate synthase reaction with the rest coming from the pyruvate dehydrogenase reaction. Under low demand conditions with increasing [Ca^2+^], the fuel utilization ratio shifts to increased fractional consumption of pyruvate, with 83±10% of acetyl-CoA derived from pyruvate at the highest [Ca^2+^] evaluated. With high ATP demand, the majority of acetyl-CoA is derived from pyruvate, regardless of the Ca^2+^ level. Our results suggest that changes in work rate alone are enough to effect a switch to carbohydrate use while in vivo the rate at which this switch happens may depend on mitochondrial calcium.

**Key Points:**

- Despite its effects on activity of mitochondrial dehydrogenases, Ca^2+^ does not substantially alter mitochondrial ATP synthesis in vitro under physiological substrate conditions. Nor does is appear to play an important role in respiratory control in vivo in the myocardium.
- We hypothesize that Ca^2+^ plays a role mediating the switch in fuel selection to increasing carbohydrate oxidation and decreasing fatty acid oxidation with increasing work rate.
- To determine if and how Ca^2+^ affects the relative use of carbohydrates versus fatty acids in vitro we measured oxygen consumption and TCA cycle intermediate concentrations in suspensions of purified rat ventricular mitochondria with carbohydrate, fatty acid, and mixed substrates at various [Ca^2+^] and ATP demand rates.
- Our results suggest that changes in work rate alone are enough to effect a switch to carbohydrate use in vitro while in vivo the rate at which this switch happens may depend on mitochondrial calcium.

## Introduction

The kinetic function of several key enzymes involved in energy metabolism is affected by calcium-dependent processes. These include cytosolic enzymes, such as phosphofructokinase (PFK), for which calcium-calmodulin-dependent oligomerization of the enzyme affects catalytic activity [1, 2], as well as several dehydrogenases present in the mitochondrial matrix [3-5]. Potentially important sites of Ca^2+^-mediated stimulation of matrix dehydrogenases are illustrated in Fig. 1. It has been proposed that in the myocardium *in vivo*, ATP, ADP, and inorganic phosphate (Pi) levels are maintained at essentially constant levels across different cardiac work rates because changes in ATP consumption rates are balanced by calcium-dependent changes in ATP production rate [3, 6]. This hypothesis, that ATP supply is matched to ATP demand based on an open-loop control system (mediated via mitochondrial Ca^2+^) lacking a closed-loop feedback control mechanism, is broadly invoked [4, 6]. But this open-loop Ca^2+^ activation hypothesis has some critical shortcomings. The first concern is that open-loop control systems are inherently unstable to environmental changes/external perturbations. For open-loop stimulation (such as via Ca^2+^) to be the sole mechanism controlling myocardial ATP production, the relationship between ATP utilization rate and the stimulatory signal (i.e., mitochondrial Ca^2+^) would have to remain exactly invariant under all physiological conditions. Otherwise, since the heart turns over its total ATP pool several times per minute, even a slight mismatch between supply and demand would be disastrous.

**Figure 1:**
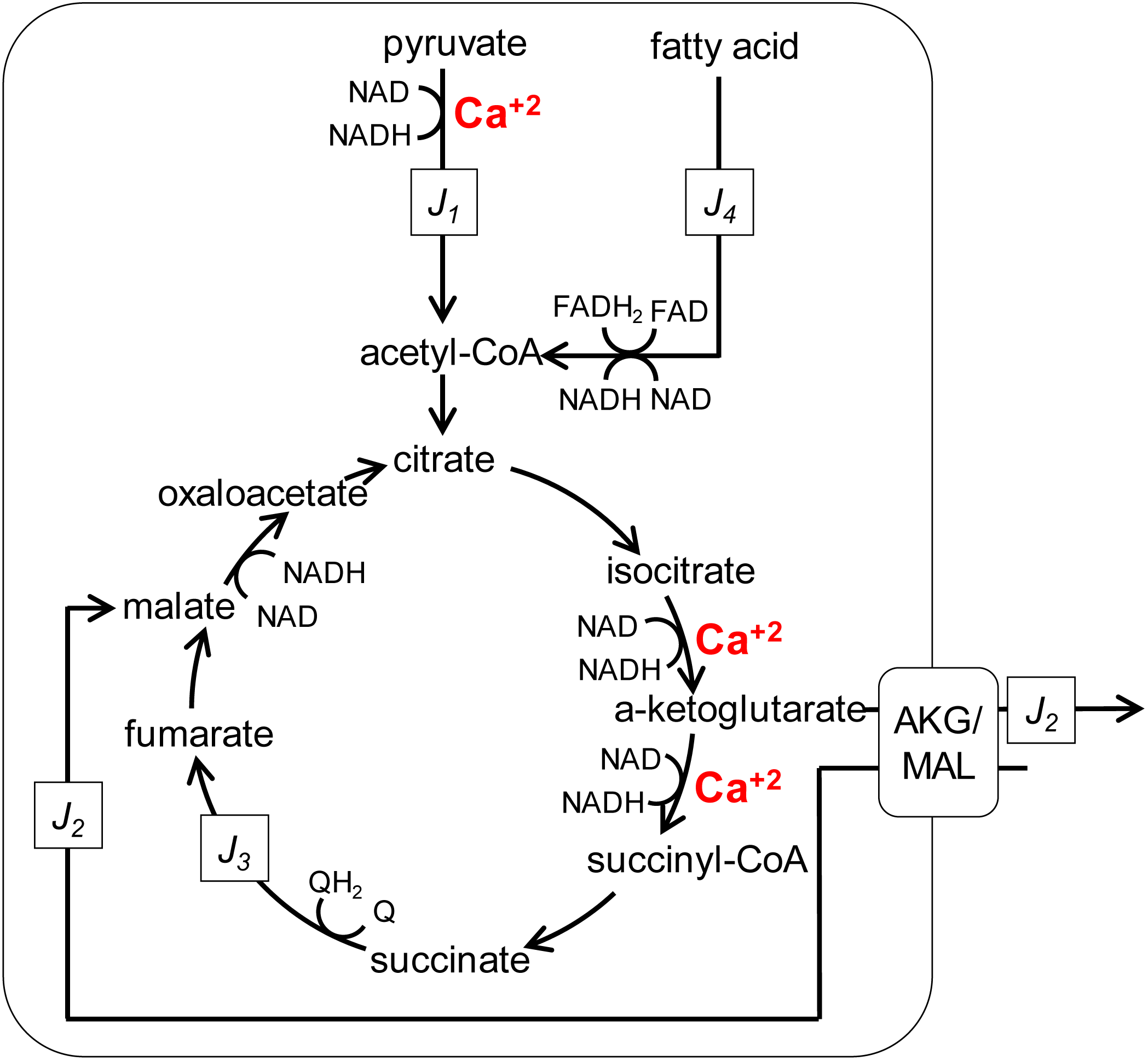
Quasi-steady-state TCA cycle fluxes. Definitions of steady-state fluxes used in the quasi-steady analysis of Eqs. (1)-(5) are illustrated. The major assumptions are that rates of change bulk concentrations of citrate, isocitrate, succinyl-CoA, succinate, fumarate, and oxaloacetate are much smaller in magnitude than rates of change of pyruvate, fatty acid, α-ketoglutarate, and malate.

Thus, some degree of closed-loop feedback control is needed to maintain cellular ATP concentration at different levels of ATP hydrolysis. It is impossible for open-loop Ca^2+^ hypothesis—on its own—to explain respiratory control *in vivo*. The alternative to the open-loop Ca^2+^ hypothesis is that respiratory control exerted by feedback of ATP hydrolysis products, introduced by Chance and Williams, concluding that “ADP, and not the inorganic phosphate level, controls the respiration rate [7].” This conclusion was later supported by *in vivo* measurements on skeletal muscle in the 1980’s in the Chance lab [8, 9]. Numerous experimental and theoretical studies have supported the feedback hypothesis in the context of skeletal muscle (e.g., [10-13]). Similarly, in the heart the concentrations of ATP hydrolysis products increase with ATP demand in the myocardium *in vivo* [14-22], in agreement with the feedback hypothesis and providing additional evidence against the open-loop Ca^2+^ hypothesis. Furthermore, our analyses suggest that a key difference between how oxidative ATP synthesis is controlled in skeletal versus cardiac muscle is that in the heart inorganic phosphate, and not ADP, controls the respiration rate [19-21, 23-25].

The third shortcoming of the open-loop Ca^2+^ hypothesis is that, while Ca^2+^ can be shown to stimulate respiration *in vitro* under certain conditions, under physiological substrate conditions the effects of Ca^2+^ on oxidative ATP synthesis *in vitro* are miniscule to modest. Panov and Scaduto [26] showed that with pyruvate and acetyl-carnitine as substrates, Ca^2+^ stimulates a maximal 3-8% increase in the apparent V_max_ and a 10-18% reduction in the apparent K_m_ for ADP for ATP synthesis. Wan et al. [27] reported a similarly small magnitude of effect of Ca^2+^ with pyruvate as substrate. Vinnakota et al. [28, 29] reported increases in the apparent V_max_ of oxidative phosphorylation of cardiac mitochondria of approximately 15-25% when external free [Ca^2+^] was raised from 0 to 350 nM. These effects are smallcompared to the 400-500% increase in ATP synthesis associated with the transition from resting to maximal work in the heart *in vivo* [19]. While the open-loop Ca^2+^ mechanism may be at work *in vivo*, its role in stimulating ATP synthesis *in vivo* may be minor compared to other mechanisms that must be at work [25].

Yet there is a growing evidence that normal mitochondrial calcium handling is required for normal cardiac mitochondrial metabolic function. Inducible fuctional knockouts of cardiac mitochondrial calcium uniporter have demonstrated that inhibition of calcium uptake has an inhibitory effect on ATP production in acute stress responses [30, 31]. Furthermore, inducible functional knockouts of cardiac mitochondrial calcium uniporter in skeletal muscle have shown to cause a metabolic shift towards fatty acid oxidation [32].

If Ca^2+^ does not play a major role in matching the steady-state oxidative ATP synthesis rate to ATP demand in the heart *in vivo*, what might be the physiological function of the Ca^2+^ sensitivity of pyruvate dehydrogenase, isocitrate dehydrogenase, and α-ketoglutarate dehydrogenase? Based on the uniporter knockout studies, we hypothesize that Ca^2+^-mediated effects on these enzymes are involved in substrate selection. Under resting conditions (in the fasted state), roughly 60% of acetyl-CoA supply to the tricarboxylic acid cycle (TCA) in the heart is derived from β-oxidation of fatty acids, with the remaining 40% derived from carbohydrate sources [33]. During exercise (high ATP demand) the ratio flips, with the majority of acetyl-CoA supplied from oxidation of carbohydrates [33, 34]. This phenomenon of increasing relative contribution from carbohydrate oxidation with increasing ATP synthesis rate is recapitulated *in vitro* in isolated mitochondria from skeletal muscle [35].

The goals of this study are to: (1.) determine if, similar to what is observed with skeletal muscle mitochondria, changes in ATP synthesis rate cause changes in fuel selection *in vitro* in suspensions of mammalian cardiac mitochondria; (2.) determine if changes in Ca^2+^ concentration cause changes in fuel selection *in vitro*; (3.) determine if and how work load and calcium independently and/or dependently influence mitochondrial substrate selection *in vitro*; and (4.) test the hypothesis that increases in mitochondrial Ca^2+^ contribute to a shift to using relatively more carbohydrate and relatively less fatty acid substrate to fuel oxidative ATP synthesis in cardiac mitochondria as ATP synthesis rate is increased. To achieve these goals, we measured oxygen consumption and bulk metabolite concentration changes during leak state (state 2 respiration) and during active oxidative phosphorylation (OXPHOS), with carbohydrate, fatty acid, and mixed (carbohydrate + fatty acid) substrates in suspensions of purified rat ventricular mitochondria. Data were analyzed using a quasi-steady-state mass balance approach to estimate TCA cycle fluxes and fractional fuel utilization under different calcium conditions and at different rates of ATP synthesis.

We find that under mixed substrate conditions fuel utilization is modulated in cardiac mitochondria *in vitro* both by ATP synthesis/demand rate and by Ca^2+^ concentration. At nominally zero [Ca^2+^], under leak state, and low ATP demand conditions, approximately 50% of acetyl-CoA is supplied to the TCA cycle from pyruvate oxidation, with the rest supplied via β-oxidation. When Ca^2+^ is added to the buffer, the fractional contribution from pyruvate increases to roughly 80% or more under low ATP demand conditions. The estimated fuel utilization ratio is also shown to shift from fatty acids to carbohydrates with increasing ATP synthesis/demand rate. With the maximal [Ca^2+^] tested, the fractional contribution from pyruvate is 55 ± 20% in the leak state, 83 ± 10% at low ATP synthesis/demand levels (corresponding to roughly 1/3 of the apparent V_max_ of oxidative phosphorylation) and 85 ± 10% at high ATP synthesis/demand levels (corresponding to roughly 1/2 of the apparent V_max_ of oxidative phosphorylation). Thus, the maximal fractional use of pyruvate is achieved at the highest Ca^2+^ level and ATP synthesis/demand rate.

## Results

Oxygen consumption rate, and concentrations of several tricarboxylic acid (TCA) cycle metabolites were measured in suspensions of purified rat ventricular mitochondria in the leak state (with substrate present but with no ADP available as a substrate for oxidative phosphorylation), and under conditions of steady ADP infusion to establish a steady rate of ATP production. These data were used to estimate steady-state substrate oxidation and TCA cycle fluxes under different substrate conditions, different Ca^2+^ concentrations, and different rates of oxidative ATP synthesis.

### Oxygen consumption rates

Fig. 2A shows representative traces of oxygen consumption flux (*J*_*O2*_) in the leak state (*t* < 0 sec.), during low-rate ADP infusion (*I* = 78.9 nmol ADP·min^-1^·(ml)^-1^, during 0 < *t* < 190 sec.), and after stopping ADP infusion (*t* > 190 sec.). In this and all other experiments, mitochondrial were added at 0.67 units of citrate synthase (U CS) activity to 2 ml of experimental buffer. Thus, expressing the ADP infusion rate (equal to ATP synthesis rate) relative to mitochondrial citrate synthase activity, *I* = 234 nmol·min^-1^·(U CS)^-1^ for the low ADP infusion rate case. Representative traces of *J*_*O2*_ are shown for three different substrate conditions PM (pyruvate and malate at initial concentrations of 350 μM and 150 uM), PCM (palmitoyl carnitine and malate at initial concentrations of 20 μM and 150 μM), and MIX (pyruvate, palmitoyl carnitine, and malate at initial concentrations of 350 μM, 20 μM and 150 μM, respectively). Fig. 2B shows similar traces, obtained in the leak state (*t* < 0 sec.), during high rate of ADP infusion (*I* = 356 nmol·min^-1^·(U CS)^-1^, 0 < *t* < 125 sec.), and after stopping ADP infusion (*t* > 125 sec.). These two ADP infusion rates correspond to roughly 1/3 and 1/2 of the V_max_ of mitochondrial ATP synthesis *in vitro*.

**Figure 2:**
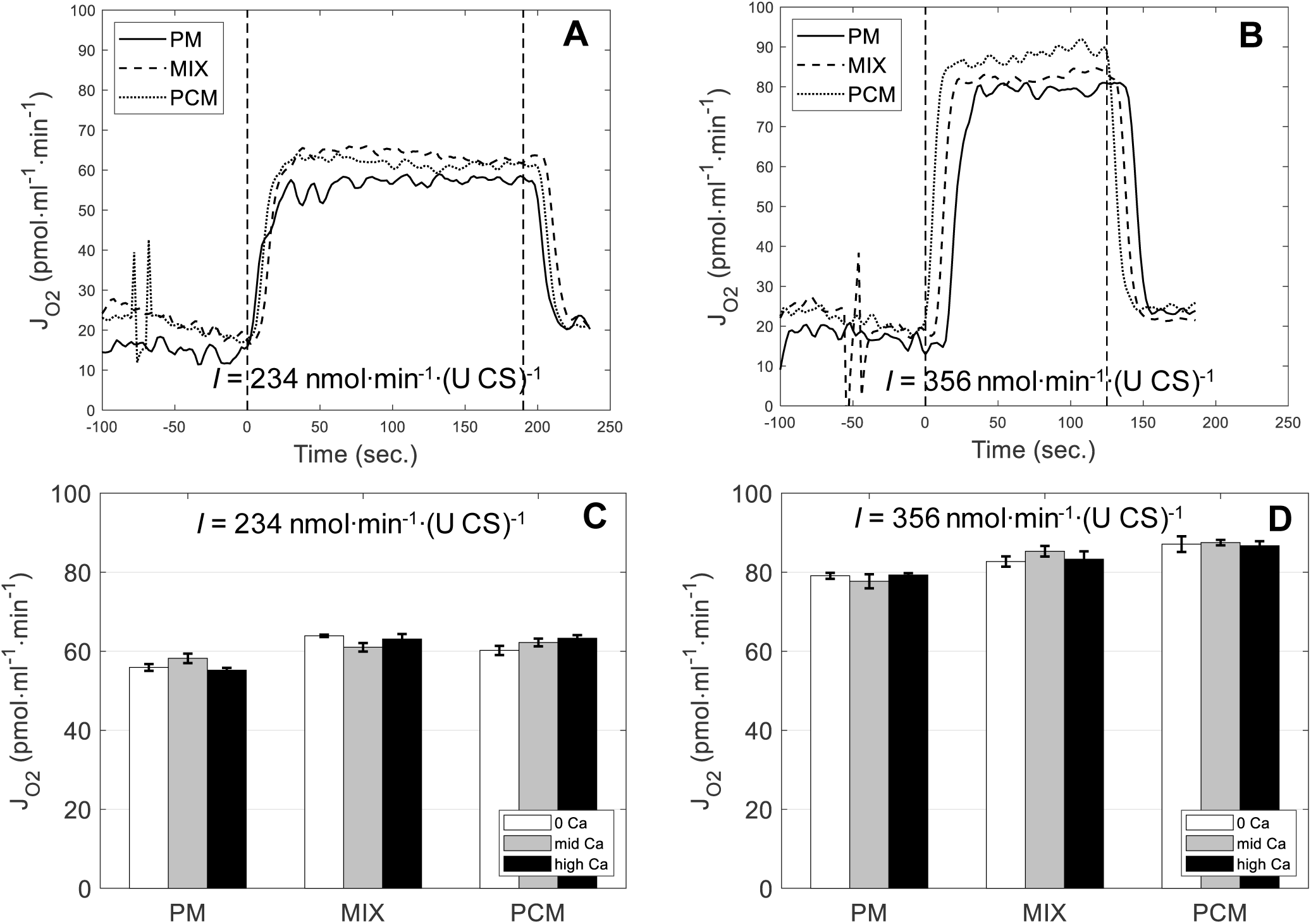
Oxygen fluxes measured by high-resolution respirometry. For all data mitochondria are suspended in a 2-ml Oroboros Oxygraph respirometry chamber at 0.337 U CS per ml. A. Time courses of oxygen consumption rate *J*_*O2*_ are shown for buffer total [Ca^2+^] = 400 μM (with 1 mM EGTA) and three different substrate conditions. Data for t < 0 correspond to the leak state. Infusion of ADP at *I* = 234 nmol min^-1^ (UCS)^-1^ begins at *t* = 0 and ends at *t* = 190 sec. B. Time courses of *J*_*O2*_ are shown for [Ca^2+^] = 400 μM and the three different substrate conditions with ADP infusion rate *I* = 356 nmol min^-1^ (UCS)^-1^. Infusion begins at *t* = 0 and ends at *t* = 125 sec. C. Data on OXPHOS state *J*_*O2*_ from N = 3 replicates are shown as Mean ± SEM for three Ca^2+^ concentrations and three different substrate conditions, and for low ATP demand (*I* = 234 nmol min^-1^ (UCS)^-1^). D. Data on OXPHOS state *J*_*O2*_ from N = 3 replicates are shown as Mean ± SEM for three Ca^2+^ concentrations and three different substrate conditions, and for high ATP demand (*I* = 356 nmol min^-1^ (UCS)^-1^). Abbreviations are PM: pyruvate + malate; MIX: pyruvate + palmitoyl-carnitine + malate; PCM: palmitoyl-carnitine + malate.

Observations from N = 4-6 biological replicates of steady-state *J*_*O2*_ during ATP synthesis (OXPHOS state) and at three different Ca^2+^ concentrations are summarized in Figs. 2C and 2D. As expected, the oxygen consumption rate *J*_*O2*_ tends to be higher when fatty the acid substrate (PCM) is oxidized compared to carbohydrate (PM). The estimated effective P/O ratios under the different calcium and substrate conditions are listed in Table 1. Here the expected trend of lower P/O ratio with PCM compared to PM is observed. Estimated P/O ratios for the MIX substrate cases tend to fall between the values estimated for PM and for PCM, although this is not the case for every combination of ATP-synthesis/ADP-infusion rate and calcium level.

**Table 1:** Summary of oxygen consumption data.

| | | Leak state<br>$J_{O_2}$ | OXPHOS I = 234 nmol min<sup>-1</sup> (UCS)<sup>-1</sup><br>$J_{O_2}$ P/O | | OXPHOS I = 356 nmol min<sup>-1</sup> (UCS)<sup>-1</sup><br>$J_{O_2}$ P/O | |
| --- | --- | --- | --- | --- | --- | --- |
| PM | [Ca<sup>2+</sup>] = 0 | 16.6± 0.28 | 55.9± 0.85 | 2.09 ± 0.03 | 79.1 ± 0.76 | 2.25 ± 0.02 |
|  | [Ca<sup>2+</sup>] = 300 μM | 17.0± 0.71 | 58.2± 1.2 | 2.01 ± 0.04 | 77.7 ± 1.78 | 2.29 ± 0.05 |
|  | [Ca<sup>2+</sup>] = 400 μM | 16.1 ± 0.21 | 55.2 ± 0.58 | 2.11 ± 0.02 | 79.3± 0.45 | 2.24 ± 0.01 |
| MIX | [Ca<sup>2+</sup>] = 0 | 22.1 ± .83 | 63.9 ± 0.3 | 1.83 ± 0.01 | 82.7 ± 1.3 | 2.15 ± 0.03 |
|  | [Ca<sup>2+</sup>] = 300 μM | 20.9 ± .75 | 61.0 ± 1.08 | 1.90 ± 0.03 | 85.3 ± 1.35 | 2.09 ± 0.03 |
|  | [Ca<sup>2+</sup>] = 400 μM | 21.0 ± .006 | 63.1 ± 1.26 | 1.85 ± 0.04 | 83.3 ± 1.99 | 2.13 ± 0.05 |
| PCM | [Ca<sup>2+</sup>] = 0 | 21.0 ± 0.76 | 60.2 ± 1.17 | 1.94 ± 0.04 | 87.1± 1.99 | 2.04 ± 0.05 |
|  | [Ca<sup>2+</sup>] = 300 μM | 20.3 ± .37 | 62.2 ± .98 | 1.88 ± 0.03 | 87.5 ± .69 | 2.03 ± 0.16 |
|  | [Ca<sup>2+</sup>] = 400 μM | 21.1 ± .56 | 63.3 ± .77 | 1.84 ± 0.02 | 86.7 ± 1.14 | 2.05 ± 0.03 |

All fluxes reported in units of nmol min<sup>-1</sup> (UCS)<sup>-1</sup>.

These measurements on their own do not provide enough information to quantify the substrate oxidation and TCA cycle fluxes under the different experimental conditions. Additional data on metabolite concentrations were obtained that, combined with the oxygen flux data, allow us to estimate quasi-steady-state fluxes.

### Metabolite concentrations

The purified mitochondria system was quenched and extracted for assay of intermediate concentrations at five different time points for the PCM and PM substrate conditions and at seven different time points for the MIX substrate condition, as indicated in the protocol detailed above. For the low ADP infusion rate experiments (*I* = 234 nmol·min^-1^·(U CS)^-1^) measurements were made at time points 70 and 110 sec. following substrate addition (during the leak state) and at 200, 240 and 280 sec for PM and PCM substrates with two additional time points at 320 and 360 sec for the MIX substrate conditions during the low ADP-infusion OXPHOS state. For the high ADP infusion rate experiments (*I* = 356 nmol·min^-1^·(U CS)^-1^) measurements were made at time points 70 and 110 sec. following substrate addition for the leak state and at 180, 210, and 240 sec. with two additional time points at 270 and 300 sec for the MIX substrate conditions during the ADP-infusion OXPHOS state. Data on pyruvate, malate and α-ketoglutarate concentrations from the PYR, PC, and MIX substrate experiments are shown in Figs. 3, 4, and 5.

**Figure 3:**
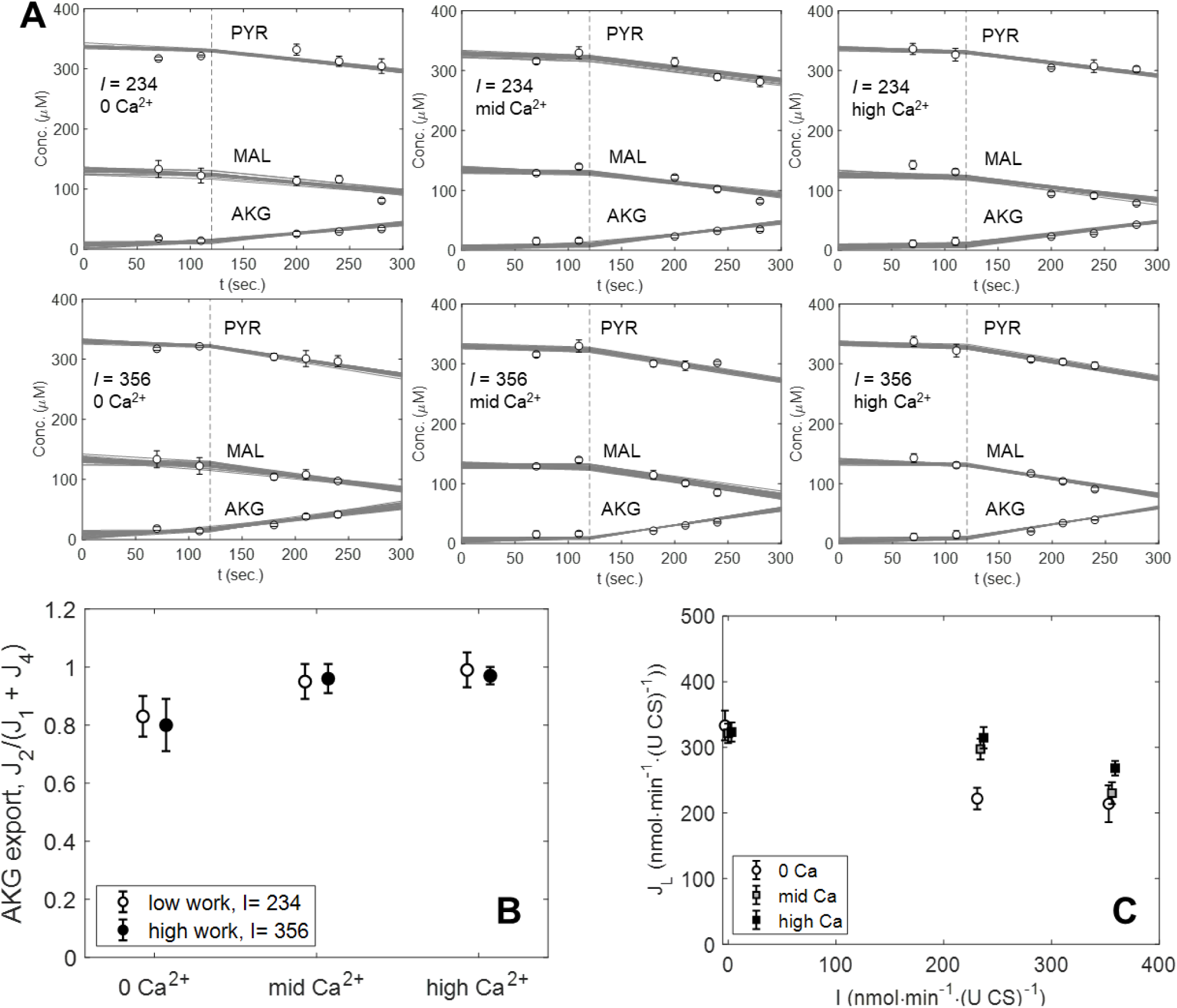
Time courses of pyruvate, malate and α-ketoglutarate during leak and OXPHOS states for PM (pyruvate + malate) substrate conditions. Straight lines are 25 Monte-Carlo samples representing uncertainty of fits of Eqs. (1) and (2) to the concentration and *J*_*O2*_ data. A. Data and model fits are shown for the [Ca^2+^] = 0, 300 μM and 400 μM conditions, with low ATP demand (*I* = 234 nmol min^-1^ (UCS)^-1^) and high ATP demand (*I* = 356 nmol min^-1^ (UCS)^-1^). B. Estimated α-ketoglutarate (J_2_/J_1_+J_4_) export for the [Ca^2+^] = 0, 300 μM and 400 μM conditions, with low ATP demand (*I* = 234 nmol min^-1^ (UCS)^-1^) and high ATP demand (*I* = 356 nmol min^-1^ (UCS)^-1^). C. Estimated leak current. The leak current *J*_*L*_ is plotted as a function of ATP demand. Estimates are obtained by combining data for all substrate conditions. (No substrate-dependent differences in *J*_*L*_ were detected.)

**Figure 4:**
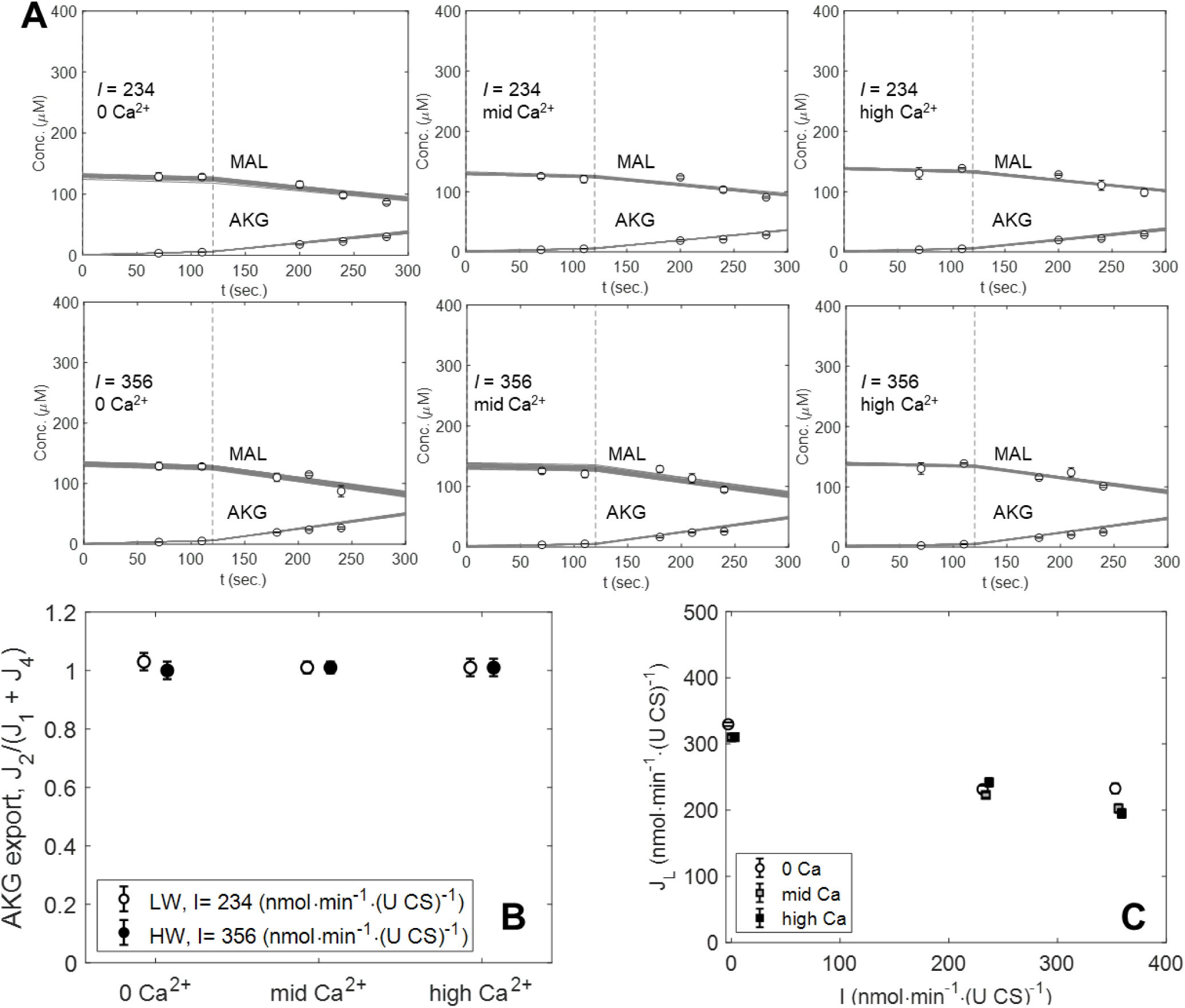
Time courses of malate and α-ketoglutarate during leak and OXPHOS states for PCM (palmitoyl-carnitine + malate) substrate conditions. Straight lines are 25 Monte-Carlo samples representing uncertainty of fits of Eqs. (3) and (4) to the concentration and *J*_*O2*_ data. A. Data and model fits are shown for the [Ca^2+^] = 0, 300 μM and 400 μM conditions, with low ATP demand (*I* = 234 nmol min^-1^ (UCS)^-1^) and high ATP demand (*I* = 356 nmol min^-1^ (UCS)^-1^). B. Estimated α-ketoglutarate (J_2_/J_1_+J_4_) export for the [Ca^2+^] = 0, 300 μM and 400 μM conditions, with low ATP demand (*I* = 234 nmol min^-1^ (UCS)^-1^) and high ATP demand (*I* = 356 nmol min^-1^ (UCS)^-1^). C. Estimated leak current. The leak current *J*_*L*_ is plotted as a function of ATP demand. Estimates are obtained by combining data for all substrate conditions. (No substrate-dependent differences in *J*_*L*_ were detected.)

**Figure 5:**
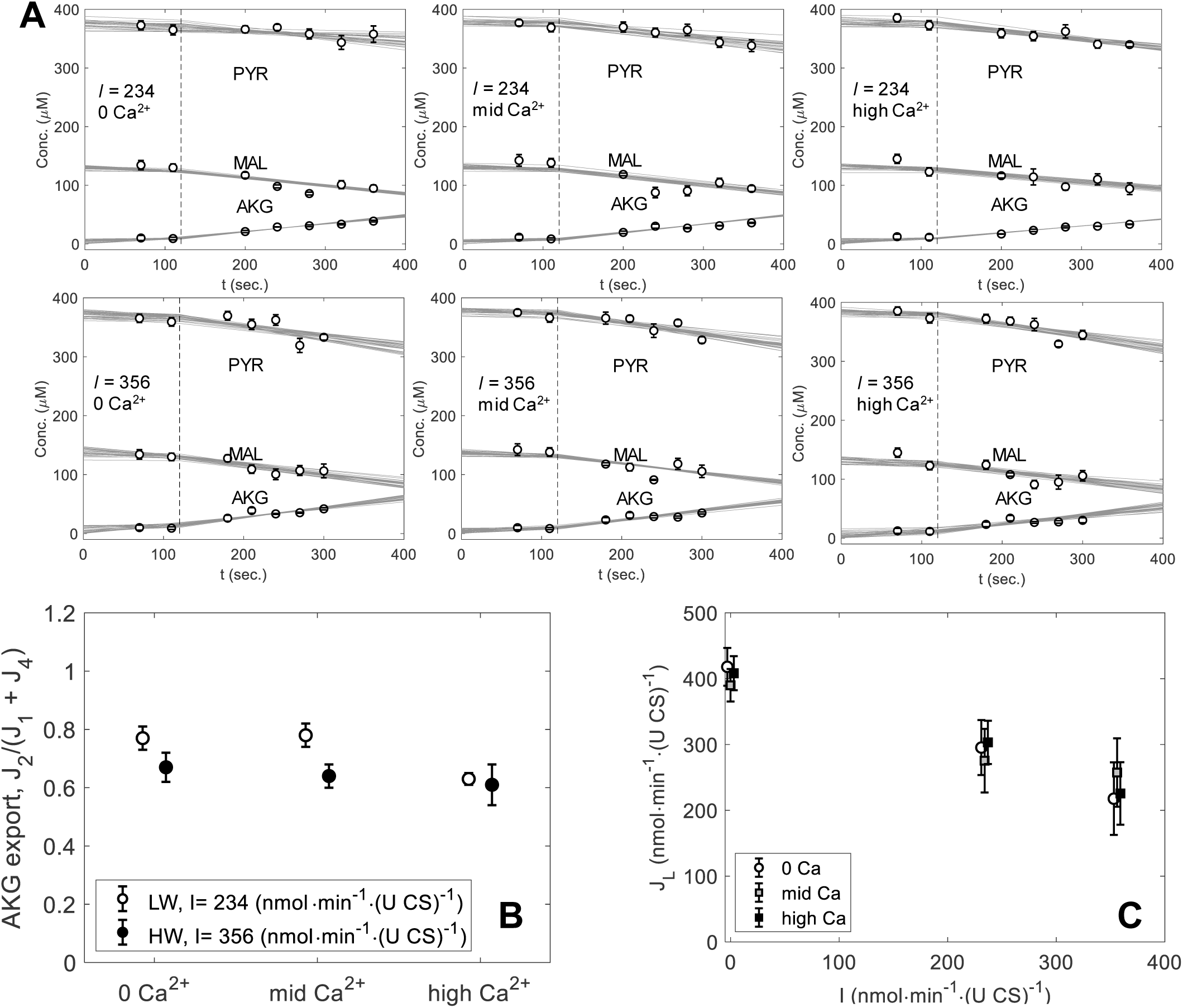
Time courses of pyruvate, malate and α-ketoglutarate during leak and OXPHOS states for MIX (pyruvate + palmitoyl-carnitine + malate) substrate conditions. A. Data and model fits are shown for the [Ca^2+^] = 0, 300 μM and 400 μM conditions, with low ATP demand (*I* = 234 nmol min^-1^ (UCS)^-1^) and high ATP demand (*I* = 356 nmol min^-1^ (UCS)^-1^). B. Estimated α-ketoglutarate (J_2_/J_1_+J_4_) export for the [Ca^2+^] = 0, 300 μM and 400 μM conditions, with low ATP demand (*I* = 234 nmol min^-1^ (UCS)^-1^) and high ATP demand (*I* = 356 nmol min^-1^ (UCS)^-1^). C. Estimated leak current. The leak current *J*_*L*_ is plotted as a function of ATP demand. Estimates are obtained by combining data for all substrate conditions. (No substrate-dependent differences in *J*_*L*_ were detected.)

Fig. 3A shows data obtained using pyruvate + malate (PM) as the substrate, at 0, 300, and 400 μM added CaCl_2_, for both ADP infusion rates. In the experimental buffer containing 1 mM EGTA, theses total calcium concentrations are associated with approximately 0, 100, and 200 nM external free [Ca^2+^]. Under these calcium conditions, with *I* = 234 nmol·min^-1^·(U CS)^-1^ roughly 40 μM of pyruvate is consumed and 25-50 μM of α-ketoglutarate is produced over the 200 second ADP infusion time period. Thus, the TCA cycle flux from citrate to α-ketoglutarate is greater than the α-ketoglutarate dehydrogenase flux. Much of the carbon entering the TCA cycle as citrate is exiting as α-ketoglutarate, rather than continuing to complete the cycle and resynthesize citrate from oxaloacetate. Instead, as has been observed previously, the exogenous malate consumed via malate dehydrogenase is the primary source of oxaloacetate for the citrate synthase reaction under these conditions [36]. A similar phenomenon is observed at the higher ADP infusion rate and under all Ca^2+^ concentrations.

Under the PCM substrate conditions (palmitoyl-carnitine + malate) no pyruvate consumption or production is observed (Fig. 4A), with α-ketoglutarate production in the same range as observed under the other substrate conditions. Under mixed substrate conditions (MIX) (Fig. 5A), the pyruvate consumption rate is much lower than observed with pyruvate + malate substrate. Although the pyruvate consumption rates are lower than in the PM condition, the α-ketoglutarate production rates are similar to those observed under the pyruvate + malate (PM) substrate conditions, indicating that a similarly low fraction of the citrate to α-ketoglutarate flux is continuing on to complete the cycle through the α-ketoglutarate dehydrogenase, succinyl-CoA synthetase, succinate dehydrogenase, and fumarase reactions.

### Quasi-steady state flux analysis

In total, oxygen consumption rate and metabolite intermediate concentrations were assayed under 27 different experimental conditions: (leak state + 2 OXPHOS states) × (3 Ca^2+^ conditions) × (3 substrate conditions). Under these different conditions, citrate, succinate, fumarate, and oxaloacetate were observed to all remain below the limit of detection (< 15 μM) of the assays employed. Based on this observation we constructed a reduced stoichiometric mass conservation model illustrated in Fig. 1, in which the fluxes *J*_*1*_ (rate of acetyl-CoA production from pyruvate dehydrogenase), *J*_*2*_ (rate of α-ketoglutarate production), *J*_*3*_ (rate of succinate dehydrogenase/complex II flux), and *J*_*4*_ (rate of acetyl-CoA production from β-oxidation) are unknowns to be estimated based on the metabolite and oxygen flux data at each experimental condition. The flux *J*_*3*_ is assumed to be equal to *J*_*1*_ + *J*_*4*_ – *J*_*2*_ under steady-state conditions. Thus, under each steady-state condition, there are three unknown steady-state fluxes, *J*_*1*_, *J*_*2*_, and *J*_*4*_ to be estimated from data on oxygen consumption rate and metabolite concentrations. We use the notation 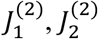, and 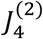 to represent fluxes in the state-2 (leak) state, and 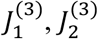, and 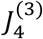 to represent fluxes in the state-3 (OXPHOS) state. In all states, the oxygen consumption flux is stoichiometrically equated to half of the rate of generation of electron donors (NADH, FADH_2_, QH_2_):

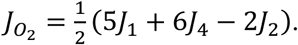

For conditions with pyruvate + malate (PM) substrate, *J*_*4*_ = 0, and we have the following equations for the leak state

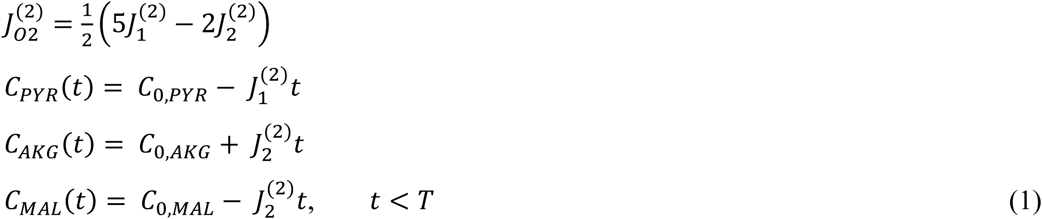

where *C*_*PYR*_(*t*), *C*_*AKG*_ (*t*), and *C*_*MAL*_(*t*) are the concentrations of pyruvate and α-ketoglutarate, *C*_0,*PYR*_ and *C*_0,*AKG*_ are the initial concentrations, and *T* is the initial time of ADP infusion. These equations assume a linear consumption of pyruvate, linear production of α-ketoglutarate, and a constant rate of oxygen consumption.

The equations for the OXPHOS state, which is initiated at *t* = *T*, are

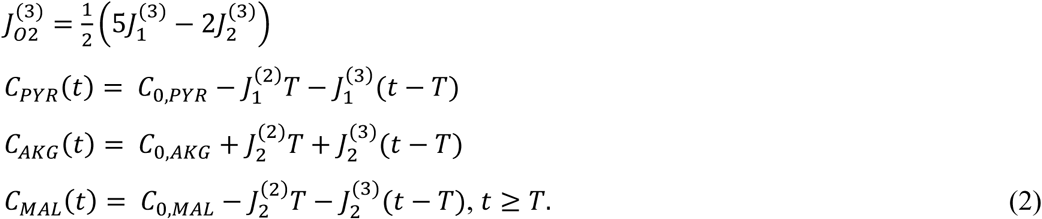

These equations assume a piecewise linear time course of pyruvate, α-ketoglutarate, and malate, as illustrated in Fig. 3A.

For experiments with PM substrate (at three different [Ca^2+^] concentrations and two different ADP infusion rates for the OXPHOS state), Equations (1) and (2) are fit to experimental data on 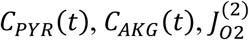, and 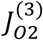 to obtain estimates of unknowns 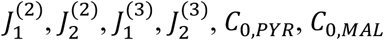, and *C*_0,*AKG*_. Even though we used initial pyruvate, α-ketoglutarate, and malate concentrations of 350, 0, and 150 μM in the experiments, respectively, we allow *C*_0,*PYR*_, *C*_0,*MAL*_, and *C*_0,*AKG*_ to be estimated in order to account for experimental variability and contamination in sample prep.

For palmitoyl-carnitine (PCM) substrate, the equations become

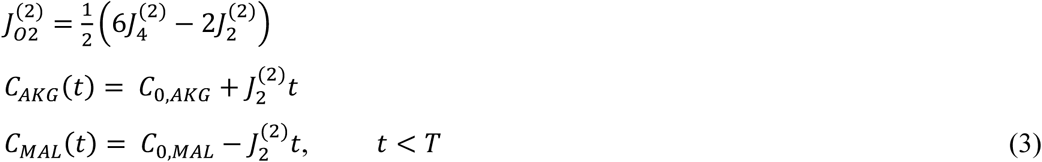

and

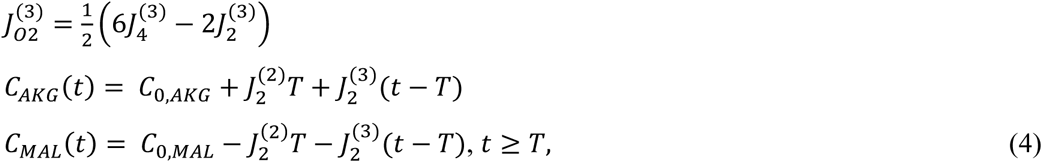

where the unknowns are 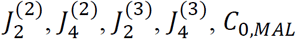, and *C*_0,*AKG*_.

With mixed substrate (MIX), the governing equations are

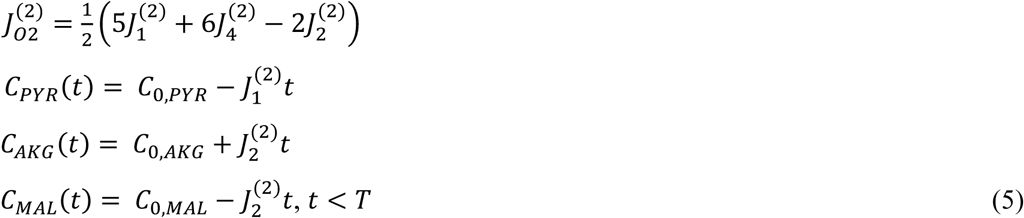

and

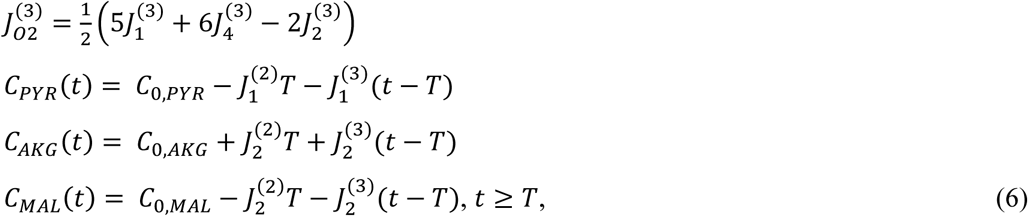

and the unknowns are 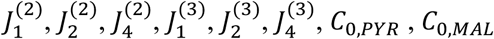, and *C*_0,*AKG*_.

For each substrate and [Ca^2+^] condition, the maximal likelihood estimates for the unknowns and for the covariances among the unknowns are obtained using the methods of Landaw and DiStefano [37]. The goodness of fit to the data and uncertainty in flux estimates are evaluated using Monte-Carlo sampling from the distributions associated with the maximally likely means and covariances for the estimated unknowns from the governing equations. For a given substrate and [Ca^2+^] condition, an individual estimate of the unknown fluxes and initial conditions yields piecewise linear fits to the pyruvate and α-ketoglutarate data and pointwise estimates of *J*_*O2*_ for the leak state and the two OXPHOS states. By drawing 10,000 samples from the flux and initial condition distributions for each substrate and [Ca^2+^] condition, we estimate the uncertainties in the estimates of the unknowns.

### Fuel utilization in vitro

Table 2 lists the estimated fluxes and uncertainties for each condition. Uncertainties are indicated as ± 1 SD confidence range. The estimates for *J*_*O2*_ in Table 2, which arise from the fits of the appropriate set of governing equations to the data, can be compared directly to the direct measurements in Table 1. The close correspondence between the measurements in Table 1 and the estimates in Table 2 (that are based on fitting both the data in Table 1 and the concentration data) provides an independent validation of the steady-state analysis.

**Table 2:**
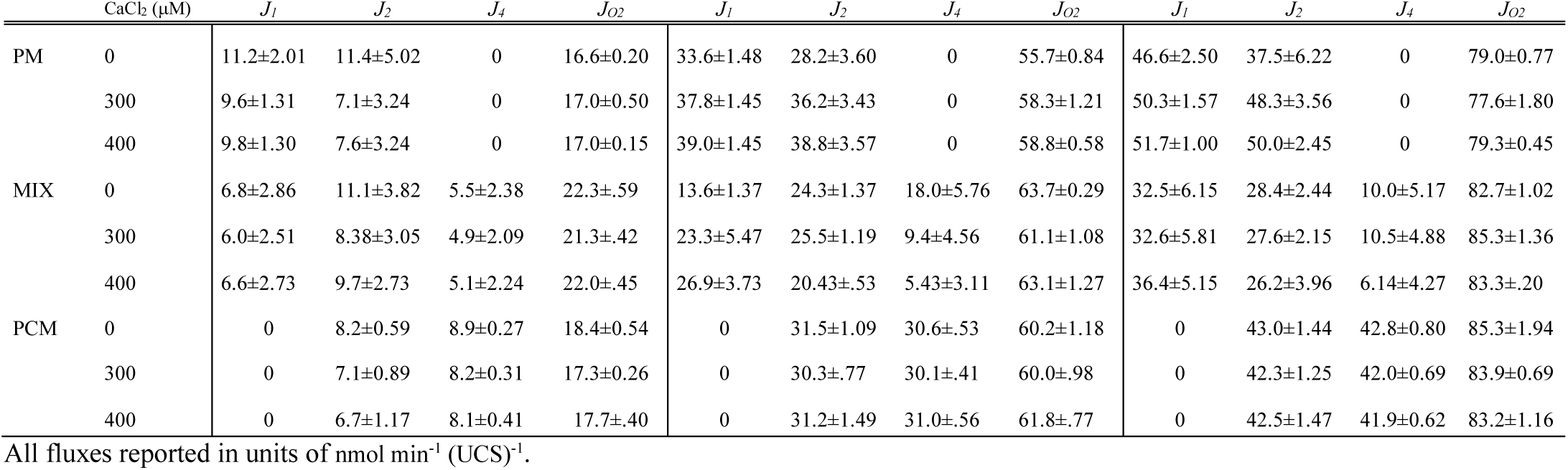
Estimated steady-state fluxes.

Representative piecewise linear fits to the concentration data are shown in Fig. 2 (PM substrate), Fig. 3 (PCM substrate), and Fig. 4 (MIX substrate) for all three [Ca^2+^] conditions. For clarity of presentation only 25 of the 10,000 independent fits are shown. The piecewise linear steady-state model effectively matches the data, and the Monte-Carlo sampling effectively matches to the uncertainty in the data, illustrated as mean ± SEM.

The flux estimates are used to estimate the fractional production of α-ketoglutarate under different OXPHOS loads and Ca^2+^ concentrations. Despite the sensitivity of α-ketoglutarate to [Ca^2+^] [38], for PM substrate the fractional export of α-ketoglutarate, *J*_2_/(*J*_1_ + *J*_4_), does not decrease with increasing [Ca^2+^] (Fig. 3B and Table 3). Analysis of the data from PCM substrate experiments yields estimates of α-ketoglutarate export that are effectively 100% of acetyl-CoA production and do not depend on ATP production/demand rate or [Ca^2+^] (Fig. 4B and Table 3). With mixed substrate (Fig. 5B and Table 3) the fractional α-ketoglutarate export is in the range of 60% to 80% and does not show a clear dependency on demand or [Ca^2+^].

**Table 3:**
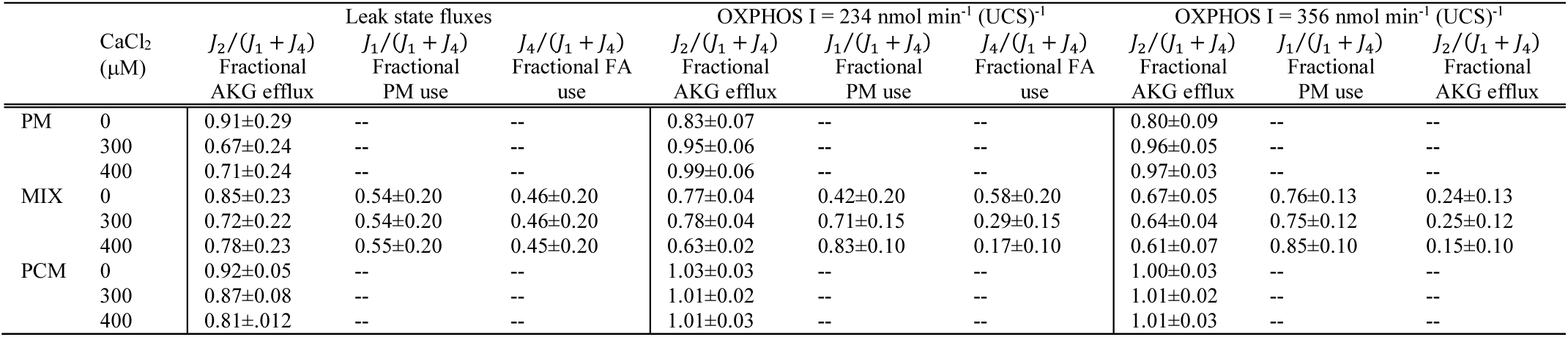
Estimated fluxes ratios.

Estimated fuel utilization (pyruvate versus fatty acid) fractions are reported in Table 3 and plotted in Fig. 6. At nominally zero [Ca^2+^], approximately 42% of acetyl-CoA is supplied to the TCA cycle from pyruvate oxidation, with the rest supplied via β-oxidation for the low ATP demand conditions. A significant increase (p = 0.033) in fractional pyruvate utilization to 83% is observed as calcium concentration increases under the low ATP demand rate condition (Fig. 6B).

**Figure 6:**
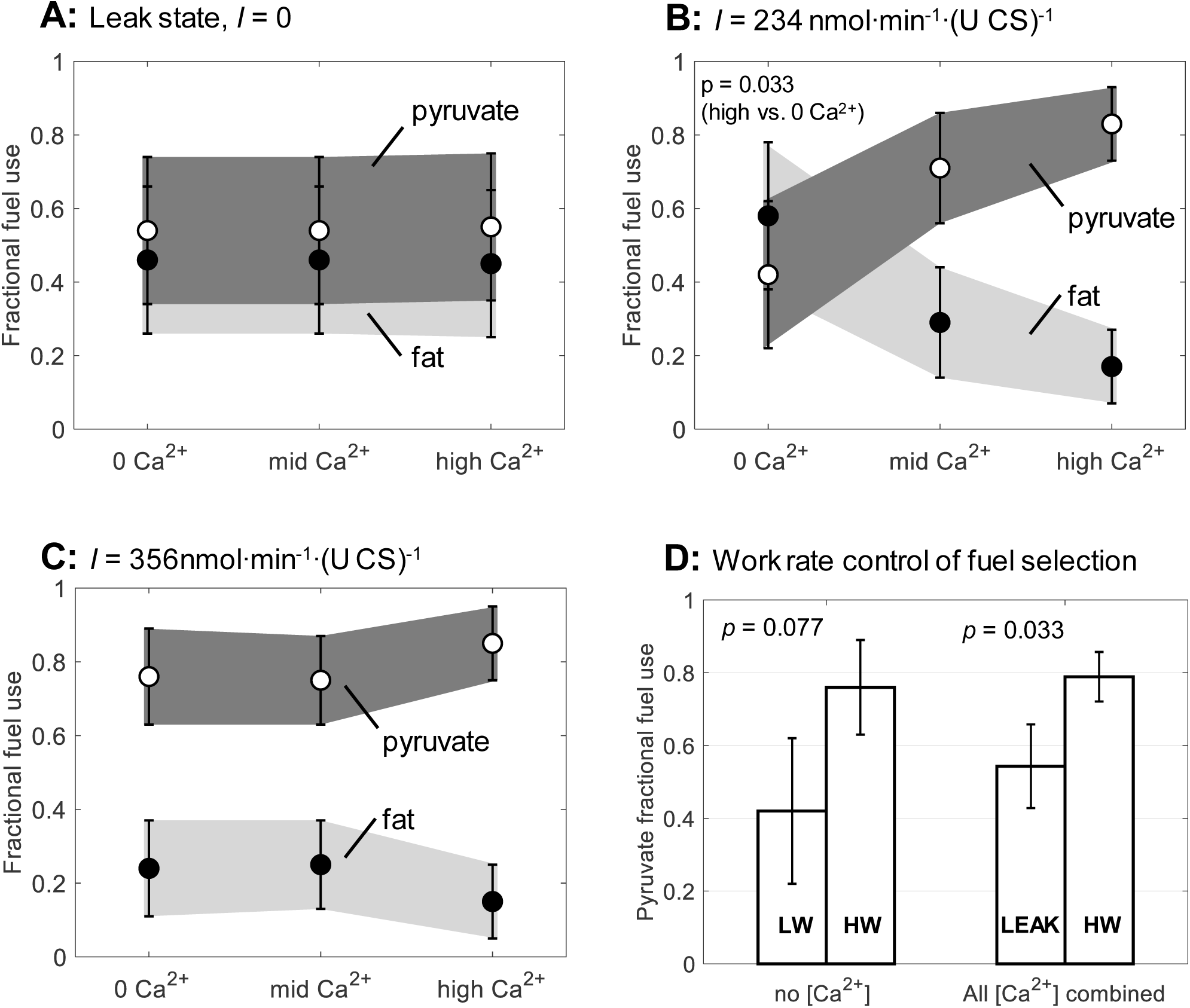
Mitochondrial fuel selection. Fractional oxidation of pyruvate versus palmitoyl-carnitine is plotted as a function of total [Ca^2+^] under (A) Leak state (*I* = 0 nmol min^-1^ (UCS)^-1^)), (B) low ATP demand (*I* = 234 nmol min^-1^ (UCS)^-1^) condition and (C) high ATP demand (*I* = 356 nmol min^-1^ (UCS)^-1^) condition. The observed trend of increasing fractional utilization of pyruvate with increasing [Ca^2+^] is statistically significant in the low ATP demand state, with estimated p = 0.033 when comparing the highest calcium vs. no calcium present (B). D. Work rate control of mitochondrial fuel selection. The observed increase in fractional utilization of pyruvate with work rate at zero Ca^2+^ is not statistically significant, with estimated p = 0.077 when comparing low vs high ATP demand. When data from all calcium concentrations are combined, an observed increase fractional utilization of pyruvate is found to be statistically significant when comparing the leak state vs high ATP demand conditions, with estimated p = 0.014.

Under high demand conditions (Fig. 6C) the estimated fractional utilization of pyruvate is 76% at zero Ca^2+^. Under this condition, addition of calcium causes a small but not statistically significant increase in the fractional contribution from pyruvate to 85 ± 10%. These results indicate that nearly all of the acetyl-CoA is being produced by pyruvate dehydrogenase in steady-state conditions under high-demand and/or high [Ca^2+^]. The ATP demand dependency is summarized in Fig. 6D. With zero added Ca^2+^ the difference in fractional fuel utilization between the low (LW) and high (HW) demand conditions of 42% versus 76% is not statistically significant, with estimated p = 0.077. Since the uncertainty on the estimates for the leak states is relatively high, statistically significant comparison between leak state and other conditions are possible only by combining all of the calcium conditions together. With all calcium conditions combined, the fractional pyruvate utilization for the leak state is 54 ± 12% and 79 ± 7% for the HW OXPHOS state (p = 0.014).

We conclude that both ATP demand and [Ca^2+^] can independently influence the fractional fuel utilization.

### Estimated leak current

The contributions of leak current to the overall oxidative flux may be estimated for the various experimental conditions probed based on the flux estimates reported in Table 2 using the following equation that estimates leak current (*J*_*L*_) as the difference between the total rate of charge pumping via the respiratory chain and the rate of charge consumption for ATP synthesis:

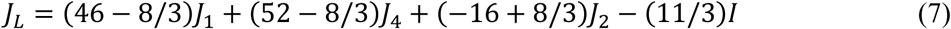

This expression is based on the following stoichiometric assumptions: (1.) each Complex I donor (NADH) generated is associated with 10 charges pumped across the inner membrane; (2.) each Complex II donor (FADH_2_ and QH_2_) generated is associated with 6 charged pumped; (3.) Each ATP generated is associated with a current of 11/3 charges. The 11/3 stoichiometric coefficient for ATP generation assumes an 8/3 stoichiometry for the ATPase [39], and 1 charge transferred by the adenine nucleotide translocase. A derivation of the expression for Eq. 7 is provided in the appendix.

Values of *J*_*L*_ estimated using these assumptions are plotted for the three different substrate conditions in Figs. 3, 4 and 5. This current is expected to be primarily composed of H^+^ ions, with secondary contributions associated with calcium cycling and other ionic currents. In the leak state (*I* = 0), the estimated leak currents are approximately 400 nmol·min^-1^·(U CS)^-1^. Since slightly fewer than 20 charges are pumped by the respiratory complexes per O_2_ molecule consumed, the leak current in the leak state may be alternatively estimated as slightly less than 20 times *J*_*O2*_ under pyruvate substrate conditions. Indeed, 20 times the average leak state *J*_*O2*_ (from Table 1) is 391 nmol·min^-1^·(U CS)^-1^. As demand is increased (*I* = 234 and 356 nmol·min^-1^·(U CS)^-1^), the estimated leak current decreases, presumably due to a decrease in magnitude of the mitochondrial membrane potential. Under OXPHOS conditions *J*_*L*_ is estimated to be approximately 200-300 nmol·min^-1^·(U CS)^-1^ for all substrate conditions. Since the ATP-producing current is 11/3 times *I*, the ratios of ATP-producing current to leak current are approximately 3:1 at *I* = 1/3 of V_max_, and 7:1 at *I* = 1/2 of V_max_.

## Discussion

This study was designed to determine if and how fuel selection *in vitro* in cardiac mitochondria is affected by ATP demand and Ca^2+^ concentration, and to test the specific hypothesis that increases in mitochondrial Ca^2+^ can cause a shift to using relatively more carbohydrate and relatively less fatty acid substrate to fuel oxidative ATP synthesis in cardiac mitochondria.

Our results demonstrate that increasing calcium concentration does effect a switch in fuel utilization under mixed substrate conditions from primarily fatty acid at nominally zero calcium to primarily pyruvate with non-zero added calcium. Furthermore, our results demonstrate that this calcium dependent fuel selection switch is also dependent on ATP demand. At the highest added calcium employed, the relative contribution of pyruvate to oxidative metabolism in mixed substrate conditions increases from 55 ± 20% at the leak state (ADP infusion rate of *I* = 0 nmol·min^-1^·(U CS)^-1^ to 83 ± 10% at an ADP infusion rate of *I* = 234 nmol·min^-1^·(U CS)^-1^ (approximately 1/3 of V_max_) to 85 ± 10% at *I* = 356 nmol·min^-1^·(U CS)^-1^ (approximately 1/2 of V_max_). These findings are analogous to the trend of increasing fraction of pyruvate utilization with increasing demand observed by Kuzmiak-Glancy and Willis [35] in skeletal muscle mitochondrial from rat and sparrow.

The trend of increasing relative contribution of pyruvate with increasing Ca^2+^ is apparent at the moderate ATP demand level. But as the ATP demand level increases from moderate to high, the fuel selection switch towards pyruvate utilization regardless of the calcium concentration. Thus, the effects of Ca^2+^ and increasing ATP demand on substrate switching appear to be additive. Furthermore, our data suggest that, at least under the *in vitro* conditions probed in this study, the primarily regulator of Ca^2+^-dependent fuel selection is pyruvate dehydrogenase.

Fluxes through downstream dehydrogenases do not show clear dependencies on Ca^2+^ concentration under any of the substrates employed here.

Based on these observations we speculate that a physiological non-zero calcium concentration is necessary for cardiac mitochondria to effectively switch from a low-capacity fuel (fats) to a high-capacity fuel (carbohydrates) in response to acute increases in workload. This mechanism may help explain observations that deletion of the cardiac mitochondrial calcium uniporter is associated with an impaired ability of the heart to acutely match ATP production to demand [30, 31].

The AKG/malate exchange has been reported to be sensitive to calcium levels in brain as well as muscle and heart mitochondria from rats [39]. But in our results, regardless of the absence or presence of calcium, the AKG and malate exchange rate remained consistent regardless of the substrate and work rate conditions used across our experimental design with constant α-ketogluterate efflux from 80%-100%. These results were unexpected as we hypothesized that if calcium enhances the substrate affinity of α-ketogluterate dehydrogenase [40], α-ketogluterate dehydrogenase flux would be increased as calcium concentration increased. The findings that fluxes through downstream dehydrogenases do not show strong dependencies on Ca^2+^ concentration supports a model in which the primarily regulator of Ca^2+^-dependent fuel selection is pyruvate dehydrogenase.

In summary, our results show that a fuel selection switch from fatty acids to greater fractional utilization of pyruvate occurs with increasing ATP demand rate and calcium in cardiac mitochondria *in vitro*. These results mirror previous observation obtained using isolated skeletal muscle mitochondria [35]. Finally, these observations are consistent with our hypothesis that calcium contributes to modulating the switch from fatty acid to carbohydrate oxidation with increasing ATP demand in vivo. This interpretation is consistent with the observations that mitochondrial calcium import is important in maintaining ATP production in acute stress responses [30, 31]. Namely, the steady-state results indicate that changes in work rate alone are enough to effect a switch to carbohydrate use. *In vivo*, the rate at which this switch happens may depend on mitochondrial calcium.

## Experimental Procedures

### Isolation of Mitochondria

Cardiac mitochondria were isolated from Male Wistar rats of 300–400 g using protocols that were approved by the Animal Care Committee of the University of Michigan. The rats were anesthetized with an intraperitoneal injection of appropriate amount of ketamine mixed with dexmed, followed by heparin. After the rat was in the deep plane of anesthesia, hearts were excised, the aortas cannulated and hearts perfused with ice-cold cardioplegia buffer (containing 25 mM KCl, 100 mM NaCl, 10 mM Dextrose, 25 mM MOPS, 1 mM EGTA) for a 5-minute period. The ventricles of the excised heart were then immediately placed in ice-cold isolation buffer containing 200 mM mannitol, 60 mM sucrose, 5 mM KH_2_PO_4_, 5 mM MOPS, 1 mM EGTA, and 0.1% BSA. The ventricles were minced with fine scissors for 3 minutes in a small beaker containing 300 ul of ice-cold isolation buffer to prevent drying of tissue. When the ventricle pieces were about 1 mm^3^, 10 ml of isolation buffer without BSA containing 3 units/ml protease were added to the minced tissue. The solution of minced tissue and isolation buffer without BSA and protease was then transferred to a Potter Elvehjem tissue grinder to be manually homogenized for maximum 3 minutes. After 3 minutes, 30 ml of ice-cold isolation buffer with BSA and 20 ul of the protease inhibitor cocktail #3 (VWR cat# 80053-854) was added to the homogenate to a final volume of 40 ml and centrifuged twice at 8000 g for 10 min at 4°C to remove the protease. The supernatant was discarded, and the pellet was resuspended in the isolation buffer to 25 ml and centrifuged at 700 g for 10 min. The pellet was then removed and the supernatant enriched with mitochondria was spun at 8000 g for 10 min. The pellet, representing the mitochondrial-enriched fraction, was resuspended in 0.2 ml of isolation buffer. To determine the content of intact mitochondria, citrate synthase (CS) activity was assessed as a quantitative marker of the mitochondrial matrix. The enzymatic activity of citrate synthase was assayed following the protocol of Eigentler et al. [40].

### Respiratory Control Index

The functional integrity of the mitochondria was determined by means of the respiratory control index defined as the maximum rate of oxygen consumption in state-3 oxidative phosphorylation (OXPHOS) divided by the state-2 (leak) state oxygen consumption rate. Both maximal OXPHOS and leak states were assessed with 5 mM pyruvate and 1 mM malate as substrates at 37° C. The OXPHOS state was assessed with ADP added at initial concentration of 500 μM. Oxygen consumption was determined at 37° C using a high resolution respirometer (Oxygraph-2K, OROBOROS Instruments Gmbh, Innsbruck, Austria). Mitochondria preparations with respiratory control index greater than 8 were considered to be of acceptable quality for our experiment.

### Steady ATP Synthesis Experiments

Purified mitochondria were resuspended in the respiration buffer (90 mM KCl, 1 mM EDTA, 5 mM KH_2_PO_4_, 50 mM MOPS, 0.1% BSA pH 7.4). The experiments were conducted with three different substrate combinations: pyruvate + malate (PYR), palmitoyl carnitine (PC) + malate, and pyruvate + PC + malate (MIX) with three different CaCl_2_ concentrations, and two different ADP infusion rates at 37° C and 7.2 pH. The final substrate concentrations were: malate: 150 uM), pyruvate: 350 μM, PC: 20 μM. Total calcium (CaCl_2_) concentrations were: 0, 300 μM, and 400 μM.

### Mitochondrial Extraction and Metabolite Assays

To measure the accumulation/utilization of various TCA cycle intermediates (pyruvate, α-ketoglutarate, malate) at various time points, samples were quenched at five different points for the PCM and PM substrate conditions and at seven different time points for the MIX substrate condition in the steady-state protocols (two during the leak state and three to five during the ADP-infusion OXPHOS state.) Samples were quenched adding 800 μl of the experimental sample to 560 μl of 0.6 M perchloric acid (PCA). The quenched samples were than centrifuged at 15,000 rpm for 5 mins at 4° C. After the centrifugation 1200 μl of the supernatant was collected in a different tube and adjusted the pH to between 6.2 and 7.4. The samples were then centrifuged once again for 5 mins at 15000 rpm at 4° C and 1 ml of the supernatant of the centrifuged samples were collected to measure the concentration of the desired metabolites enzymatically.

For all of the 18 experimental conditions, samples were collected at 70 and 110 seconds during the leak state. For the low-workload condition, three additional samples were collected at times 200, 240, and 280 seconds for PYR and PC substrates with two more time points at 320 and 360 seconds for the MIX substrate conditions. During the high workload condition, samples were collected at 180, 210, and 240 seconds for PYR and PC substrates with two additional time points at 270 and 300 seconds for the MIX substrate conditions. For each sample collection time, at least 4 (6 replicates in the mixed sample) replicates were obtained.

### Metabolite Assays

Metabolite assays were adapted from the methods of Williamson and Corkey [41]. For all measured metabolites NADH-linked assays were used, where the oxidation/reduction of specific metabolites are linked changes in NADH, monitored by changes in absorbance at 340 nm. Assays were run using neutralized extract in a final assay volume of 1 ml. Standard curves were obtained daily.

The α-ketoglutarate (AKG) concentration was measured by coupling glutamate oxaloacetate transaminase (GOT, EC: 2.6.1.1) with malate dehydrogenase (MDH, EC: 1.1.1.37) and adding excess of aspartate (ASP) to the system and obtaining the difference between NADH concentration before and after adding ASP:

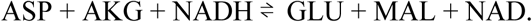

Pyruvate (PYR) concentration was measured by adding lactate dehydrogenase (LDH, EC: 1.1.1.27) to catalyze PYR reduction and NADH oxidation and by computing the difference in NADH before and after adding LDH:

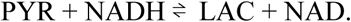

The enzyme malate dehydrogenase (MDH, EC: 1.1.1.37) catalyzes the conversion of Malate (MAL) to oxaloacetate (OAA). Coupling the reaction with glutamate oxaloacetate transaminase (GOT, EC: 2.6.1.1), Malate concentration was estimated by computing the difference between NADH concentration before and after adding GOT and MDH:

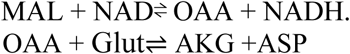

## Competing Interests

The authors declare that they have no competing interests.

## Author Contributions

EJ, RKD and DAB conceived and designed the research. EJ, SK, SKD, NN, FVdB, and AH performed the experiments. EJ, DAB, and HSC analyzed the data. EJ and DAB interpreted the results. EJ prepared the figures and drafted the manuscript. All authors edited and revised the manuscript. All authors approved the final version of manuscript. All authors agree to be accountable for all aspects of the work in ensuring that questions related to the accuracy or integrity of any part of the work are appropriately investigated and resolved. All persons designated as authors qualify for authorship, and all those who qualify for authorship are listed.

## Acknowledgments

This work supported by NIH grants HL144657, HL122199 and T32GM008322.

## Appendix

The leak current (*J*_*L*_) is calculated as the difference between the total rate of charge pumping via the respiratory chain and the rate of charge consumption for ATP synthesis. The rate of charge pumping via the respiratory chain is computed from the rates of generation electron NADH, FADH_2_, and QH_2_ associated with the flux estimates of electron donor generating enzymes associated with fluxes *J*_*1*_, *J*_*4*_, and *J*_*2*_ (Fig. 1):

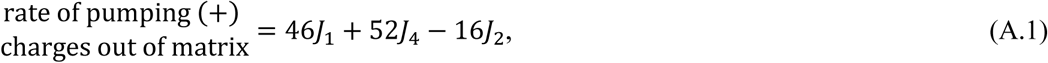

assuming that each Complex I donor (NADH) generated is associated with 10 charges pumped across the inner membrane and each Complex II donor (FADH_2_ and QH_2_) generated is associated with 6 charged pumped.

The rate of charge consumption for ATP synthesis is estimated by equating the rate of ATP synthesis I to contributions from oxidative phosphorylation and substrate-level phosphorylation by Complex II:

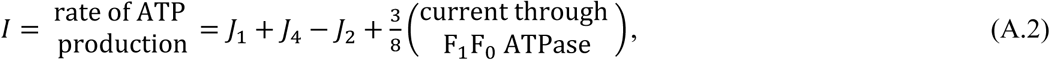

where *J*_1_ + *J*_4_ - *J*_2_ is the substrate-level phosphorylation rate, and each ATP synthesis step is associated with an average of 8/3 charges transferred in to the matrix.

The total current used for ATP synthesis is computed as sum of F_1_F_0_ ATPase current plus ANT (adenine nucleotide translocase) current:

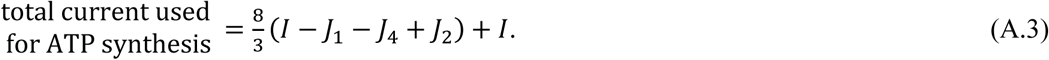

Subtracting (A.3) from (A.1), we obtain

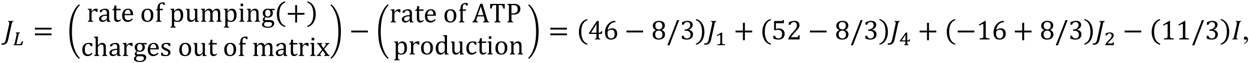

which is Equation (7).

## Notes

#### Summary of Updates

A large number of new experiments for the mixed substrate conditions were performed with additional replications, and with new data for longer time courses. These new data provided more accurate estimates of fluxes for these conditions, with lower statistical uncertainties than in the previous version. As a result that the statistical confidence in the interpretations is markedly improved compared to the previous version.

